# Social Closeness and Reward Sensitivity Enhance Corticostriatal Function during Experiences of Shared Rewards

**DOI:** 10.1101/2023.10.19.562908

**Authors:** David V. Smith, Ori Zaff, James B. Wyngaarden, Jeffrey B. Dennison, Daniel Sazhin, Jason Chein, Michael McCloskey, Lauren B. Alloy, Johanna M. Jarcho, Dominic S. Fareri

## Abstract

Although prior research has demonstrated enhanced striatal response when sharing rewards with close social connections, less is known about how individual differences affect ventral striatal (VS) activation and connectivity when experiencing rewards within social contexts. Given that self-reported reward sensitivity and level of substance use have been associated with differences in VS activation, we set out to investigate whether these factors would be independently associated with enhancements to neural reward responses within social contexts. In this pre-registered study, participants (N=45) underwent fMRI while playing a card guessing game in which correct or incorrect guesses resulted in monetary gains and losses that were shared evenly with either a close friend, stranger (confederate), or non-human partner. Consistent with our prior work, we found increased VS activation when sharing rewards with a socially close peer as opposed to an out-of-network stranger. As self-reported reward sensitivity increased, the difference in VS response to rewards shared with friends and strangers decreased. We also found enhanced connectivity between the VS and temporoparietal junction when sharing rewards with close friends as opposed to strangers. Finally, exploratory analyses revealed that as reward sensitivity and sub-clinical substance use increase, the difference in VS connectivity with the right fusiform face area increases as a function of social context. These findings demonstrate that responsivity to the context of close friends may be tied to individual reward sensitivity or sub-clinical substance use habits; together these factors may inform predictions of risk for future mental health disorders.

## 1. Introduction

From purchasing a car to getting engaged, important life decisions often are made in social contexts—whether around strangers or loved ones. Social influence can profoundly shape reward-related decision making (Dennison et al., 2022; Smith & Delgado, 2015; Powers et al., 2022), including the emergence of maladaptive choices (O’Brien et al., 2011). At the neural level, both social context and individual traits influence how rewards are experienced and valued. Specifically, the ventral striatum (VS)—a central hub for encoding subjective value (Haber & Knutson, 2010; Middleton & Strick, 2000)—responds not only to monetary rewards themselves, but also to their social context. For instance, sharing monetary gains with a friend enhances VS responses relative to sharing with strangers or nonsocial partners (Fareri et al., 2012), accompanied by parallel shifts in subjective satisfaction, excitement, and social behavior (Dziura et al., 2022). However, this “social boost” is not uniform across individuals; some show stronger neural differentiation based on social context than others, raising the question of what drives these individual differences in neural sensitivity to social reward.

Here, we propose an integrative framework in which social context, trait-level reward sensitivity, and substance-use risk jointly shape corticostriatal responses during shared reward processing. The posterior temporoparietal junction (TPJ), fusiform face area (FFA), posterior cingulate cortex (PCC) and medial prefrontal cortex (mPFC) collectively form a broader “social brain” network supporting social attention, mentalizing, self-referential processing and relational valuation (Bhanji & Delgado, 2014; Hutcherson et al., 2015; Lockwood et al., 2020; Mars et al., 2012a). A number of these regions including the mPFC and PCC are importantly implicated in encoding reward value (e.g., Bartra et al., 2013; Acikalin et al., 2017). Functional connectivity between these regions and the VS previously has been linked to generosity and enhanced reward valuation within social contexts (Haeger et al., 2014; Park et al., 2017; Tusche et al., 2016). Building on this literature, we hypothesized that shared rewards would elicit differential VS activation and VS–TPJ coupling depending on the social partner, with the strongest responses for close friends, intermediate responses for strangers, and the weakest responses for nonsocial (computer) partners—and that these neural patterns would be modulated further by key person-level traits.

A central trait of interest in this framework is reward sensitivity, defined as the degree to which appetitive stimuli motivate approach behavior (Carver & White, 1994; Kim et al., 2015; Nusslock et al., 2007). Both unusually high and unusually low reward sensitivity have been linked to mood disorders, addiction risk, and striatal dysfunction (Chat et al., 2022; Nusslock & Alloy, 2017; Volkow et al., 2010). Given these findings, we modeled both linear and quadratic relationships between reward sensitivity and corticostriatal responses. We reasoned that aberrant sensitivity at either extreme could either diminish distinctions across social partners or exaggerate neural reactivity to preferred contexts. Additionally, we examined sub-clinical substance use, another trait closely associated with altered reward processing (Franken & Muris, 2006; Kober et al., 2010; Shadur & Hussong, 2014; Strickland & Smith, 2014). We hypothesized that reward sensitivity and substance use might jointly shape individuals’ neural responses to social contexts, reflecting combined risk factors for impaired reward valuation and altered social cognition.

Despite growing interest in the neuroscience of social reward, few studies have directly examined how individual differences moderate neural responses to shared outcomes across varying social contexts. To address this gap, we employed a card-guessing task in which participants received monetary rewards shared with a close friend, a stranger, or a computer partner (Delgado et al., 2000; Fareri et al., 2012) during functional magnetic resonance imaging (fMRI). Trait reward sensitivity and substance use were examined as independent moderators of neural activation and functional connectivity during reward receipt. In our preregistered hypotheses, we predicted that social closeness (friend vs. stranger or computer) would enhance subjective reward ratings, VS activation, and VS–TPJ connectivity—and that these social context effects would be independently moderated by trait reward sensitivity and substance use.

Specifically, we anticipated that VS reactivity to sharing rewards with friends relative to less close or nonsocial partners would have a U-shaped relationship with reward sensitivity, such that we would see the highest levels of reactivity for friends (vs. others) in those with the highest and lowest levels of reward sensitivity. Such a pattern would be indicative of dysregulated reward valuation. We also expected individuals with elevated levels of substance use to exhibit increased VS activation and stronger connectivity between VS and mentalizing regions, particularly in the friend versus computer contrast. Although we initially intended to explicitly test interactions involving perceived closeness, high variability and measurement limitations of closeness ratings (Inclusion of Other in Self scale; IOS) led us to rely on the friend versus stranger contrast as a practical proxy for interpersonal closeness. These clearly articulated predictions guided our confirmatory region-of-interest analyses and exploratory whole-brain analyses, ensuring a theory-driven approach to understanding how trait-level factors modulate neural responses to socially shared reward.

## 2. Methods

### 2.1. Participants

Our preregistered goal was to collect data from 100 participants aged 18–22 (see preregistration: https://aspredicted.org/blind.php?x=SFX_MXL). However, due to constraints imposed by the COVID-19 pandemic, we were ultimately able to collect data from 52 participants who completed at least one run of the Shared Reward task as part of a broader neuroimaging session. We applied preregistered exclusion criteria to determine our final sample. Three participants were excluded for excessive head motion, defined using MRIQC quality metrics (Esteban et al., 2017) as either mean framewise displacement (fd_mean) greater than 1.5 times the upper bound of the interquartile range or temporal SNR below 1.5 times the lower bound. Two additional participants were excluded for failing to engage with the behavioral task (>20% missed responses), and two were excluded for incomplete behavioral or survey data due to technical issues. These exclusions yielded a final sample of 45 participants (mean age = 21.11 years, SD = 1.83; 36.4% male). A sensitivity analysis conducted in G*Power indicated that with 45 participants, α = .05, and power = .80, the study was able to detect an effect size of f^2^ = 0.18 or larger for a single additional predictor. While effects below this threshold could still be present and observable, this estimate helps contextualize the lower bound of reliably detectable effects.

Each participant was asked to refer a same-gendered close friend, who submitted a facial photograph used in the Shared Reward task. Participants were recruited through the Temple University Psychology and Neuroscience Department’s participant pool and through community outreach, including flyers and online advertisements. All participants were compensated for their time at a rate of $25 per hour for fMRI and $15 per hour for behavioral testing. They also received performance-based bonuses for other neuroeconomic tasks administered during the session (not reported here; see Smith et al, 2024 for details), resulting in a base payment of approximately $100 and an average bonus of $50. Participants recruited through the university pool received equivalent research credit in lieu of base compensation but remained eligible for monetary bonuses tied to task performance.

### 2.2. Procedures

Recruitment and procedural methods were approved by the Temple University IRB. Participants began the study by completing an initial interest screener. After a behavioral consent form, the screener involved completing the Behavioral Inhibition System and Behavioral Activation System Scale (BIS/BAS; Carver & White, 1994) and the Sensitivity to Punishment and Reward Questionnaire (SPSRQ; Torrubia et al., 2001) on Qualtrics. Participants who provided similar responses on both measures of reward sensitivity (i.e., within one quintile) were contacted to participate in the study (Alloy et al., 2009). Participants reporting substantially different levels of reward sensitivity (i.e., more than one quintile apart) were excluded, as such discrepancies may reflect inattentiveness, disengagement, or other factors that compromise the reliability of self-report data.

Participants were additionally excluded if they were unwilling to abstain from drinking alcohol or using recreational substances within 24 hours of the MRI scan. Those taking psychoactive medications were not recruited. Participants who passed the screener were run through a mock version of the scan to train on reducing head motion. A breathalyzer test and urine drug screen were then conducted to ensure that performance and/or brain activation at the time of the scan was not confounded by recent substance use or substances still detectable in participants’ systems. Of participants who passed the initial screener, two were excluded after testing positive for morphine and amphetamine usage. Five participants who tested positive for marijuana were included in our final sample. Participants who tested positive for marijuana were retained in accordance with our IRB-approved protocol. This decision reflects the pharmacokinetics of marijuana: unlike many other substances, its metabolites can remain detectable in urine long after acute effects have subsided. Thus, a positive marijuana screen does not indicate active intoxication or impairment at the time of scanning.

Following these procedures, participants underwent fMRI for approximately 1.5 hours, during which they completed the Shared Reward task (described below) along with three additional tasks: a monetary incentive delay task (Wyngaarden et al., 2025), a social reward task (Wyngaarden et al., 2024), and a social decision-making task (Sazhin et al., 2024). Task order was counterbalanced across participants. The Shared Reward task lasted approximately 15 minutes. After the scan, participants completed additional surveys and behavioral tasks outside the scanner. The total session, including consent, screening, scanning, and post-scan assessments, lasted approximately 2.5 hours. Full protocol details and open-access data are available via OpenNeuro, as described in our accompanying data descriptor (Smith et al., 2024).

### 2.3. Shared Reward Task

We administered a card-guessing game during fMRI to assess participants’ neural responses to monetary rewards and losses experienced in social and non-social contexts (adapted from Delgado et al., 2000; Fareri et al., 2012). On each trial, a question mark appeared on the screen, and participants had 2,500 ms to guess whether a hidden card number was higher or lower than 5 by pressing a button with their right index or middle finger. Once a response was made, the question mark turned orange and remained on screen until the trial advanced. The true card value was then revealed, along with an outcome cue: a green upward arrow for a win (correct guess), a red downward arrow for a loss (incorrect guess), or a white double-headed arrow for a neutral trial (card value = 5). If no response was made in time, a number sign appeared instead of a card, indicating a missed trial (see Fig. 1). Each trial concluded with a 750 ms fixation cross before the next decision phase. Participants completed two runs, each lasting 6 minutes and 54 seconds.

**Figure 1.**
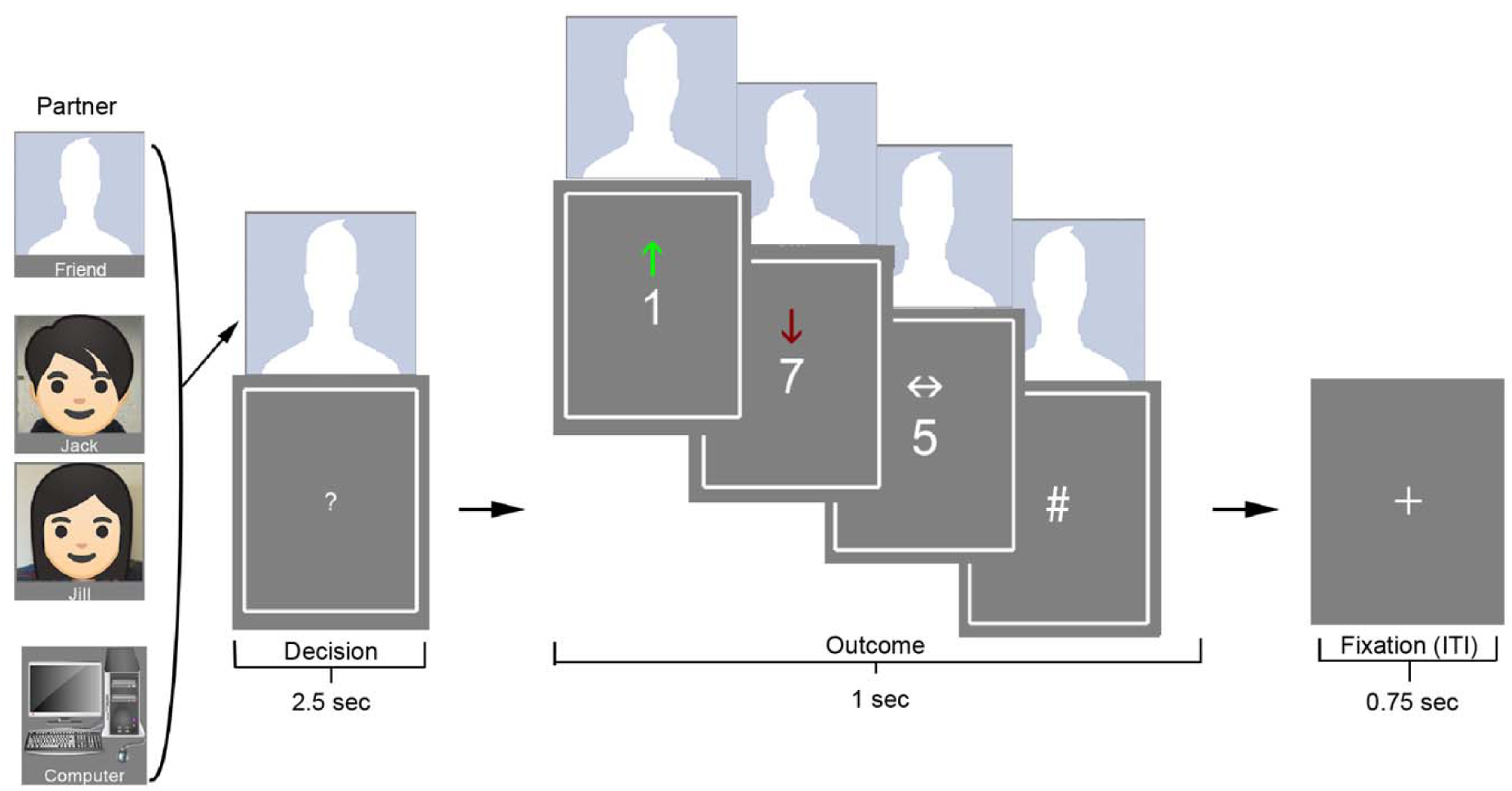
Task Structure. Participants completed a card-guessing task adapted from Delgado et al. (2000) and Fareri et al. (2012), in which they guessed whether a hidden number (1–9) was higher or lower than 5. On each trial, participants were paired with one of three partners: a computer, a same-gender confederate (Jack/Jill), or a close friend. A photo of the partner appeared above the card from the beginning of the decision phase through the outcome. Actual photos of the confederates were used in the task; the cartoon images shown here are just used for illustrative purposes. Correct guesses (green up-arrow) resulted in a $10 gain; incorrect guesses (red down-arrow) resulted in a $5 loss; and a neutral outcome (number = 5) was indicated by a white side-to-side arrow. Participants were told that earnings would be shared equally with the displayed partner on each trial.

Correct guesses earned a monetary reward of $10, whereas incorrect guesses resulted in a $5 loss. Although gain and loss magnitudes were not matched, this asymmetry reflects longstanding design precedent in the reward literature (e.g., Delgado et al., 2000) and is theoretically grounded in loss aversion frameworks, which posit that losses exert greater psychological impact than equivalent gains (Kahneman and Tversky, 1979). Each trial was paired with one of three partners: a close friend (identified and invited by the participant), a stranger (a trained confederate introduced as a past participant), or a non-human control (computer). An image of the current partner appeared above the card from the start of the decision phase through the end of the outcome phase, indicating who would share in the outcome.

Trials were organized in a blocked design, with partner identity held constant within each 8-trial block. Blocks also were grouped by outcome context—either mostly reward or mostly loss— such that 6 of 8 trials were biased toward one outcome type, with 2 trials that were neutral or of the opposite valence (cf. Barch et al., 2013). There were 18 blocks per participant, with 1–2 blocks per condition per run, consistent with validated block structures used in large-scale datasets such as the Human Connectome Project (HCP). Across subjects, the mean (SD; range) number of trials was 20.29 (7.74; 6–31) for friend reward, 20.16 (7.34; 7–30) for friend loss, 19.84 (5.87; 7–28) for stranger reward, 20.14 (5.59; 4–28) for stranger loss, 19.78 (7.28; 4–30) for computer reward, and 19.69 (7.61; 6–32) for computer loss. Block order and trial types were fully randomized across participants. Blocks were separated by a jittered fixation cross lasting 8–12 seconds to allow the hemodynamic response to return to baseline. Participants were told that one trial would be randomly selected at the end of the session and that the outcome from that trial would be shared equally with the partner (i.e., $5 added for a win or $2.50 deducted for a loss). Money shared with friends and strangers was sent to them after the study; money from computer trials was returned to a lab fund.

Immediately following the scan, participants completed a brief follow-up in which they rated the emotional salience of winning and losing with each partner (e.g., “How did it feel…?”) on a scale from –5 (very negative) to +5 (very positive). The full task and post-scan ratings were programmed in PsychoPy 3 (Peirce et al., 2019). Two participants did not complete the partner rating portion due to time constraints; analyses involving these ratings include N = 43.

### 2.4. Individual Difference Measures

#### 2.4.1. Trait Reward Sensitivity

Trait reward sensitivity was assessed using the Behavioral Activation Scale (BAS; Carver & White, 1994) and the Sensitivity to Reward subscale (SR) of the Sensitivity to Punishment and Sensitivity to Reward Questionnaire (SPSRQ; Torrubia & Tobeña, 1984). These scales measure overlapping but distinct dimensions of behavioral approach motivation and have been widely used in studies linking reward sensitivity to striatal reactivity, mood disorders, and addiction risk (Alloy et al., 2006, 2012; Kim et al., 2015). In our sample, BAS and SR scores were strongly correlated (r = .68), supporting their combination into a single index. To do so, we z-scored each measure and summed the scores, preserving individual variability while placing both scales on a common metric.

Given evidence that both unusually low and unusually high reward sensitivity are associated with maladaptive outcomes (e.g., Alloy et al., 2009; Bart et al., 2021; Franken & Muris, 2006), we modeled both linear and quadratic effects of the composite score. To reduce the undue influence of extreme values when computing the quadratic term, we binned the composite into deciles and then squared and mean-centered the resulting values. This transformation emphasizes departures from normative levels of reward sensitivity while minimizing the risk that outliers drive second-order effects (Büchel et al., 1998). Although this binning approach was not preregistered, it represents a principled refinement consistent with our theoretical rationale and analytic goals.

#### 2.4.2. Subclinical Substance Use

Substance use was operationalized as a composite score consisting of the sum of z-scores for the Alcohol Use Disorders Identification Test (AUDIT; Saunders et al., 1993) and the Drug Use Disorders Identification Test (DUDIT; Berman et al., 2002). The AUDIT is a 10-item self-report measure assessing alcohol consumption and related problems. The DUDIT is an 11-item self-report measure assessing frequency and disruptiveness of non-alcoholic drug use, referencing substances such as marijuana, amphetamines, and others. We summed the z-scores to reflect overall substance-use risk across both alcohol and drugs, consistent with our hypotheses and the dimensional nature of use in this population. Whereas the AUDIT and DUDIT capture distinct behaviors, their conceptual overlap and shared purpose as substance-use screeners support their integration into a single composite. We also binned the composite scores into deciles to reduce the influence of extreme values and to parallel our approach to reward sensitivity.

Given that participants were screened to exclude those unable to abstain from substance use, we consider this a sub-clinical sample. Both scales include thresholds for clinically significant impairment. AUDIT scores of 1–7 are considered low-risk, 8–14 indicate hazardous or harmful use, and 15+ reflect probable dependence. Scores in our sample ranged from 0 to 14 (M = 3.6, SD = 3.5). For the DUDIT, scores above 24 suggest dependence, while lower thresholds (e.g., >1 for women, >5 for men) indicate problematic use (Basedow et al., 2021). DUDIT scores ranged from 0 to 28 (M = 2.3, SD = 5.8). Two participants exceeded the dependence threshold on the DUDIT, but both had low AUDIT scores. Overall, we characterized the sample as sub-clinical. Of the 45 participants included in the analyses, eight abstained from alcohol use. Twenty-one participants reported no use of alcohol or other substances. Among those who endorsed drug use, the most commonly reported substances were Ritalin/amphetamines (n = 14), marijuana (n = 2), cocaine (n = 1), and solvents/inhalants (n = 2); the latter three also reported stimulant use.

### 2.5. Behavioral Analyses

Behavioral measures were assessed in accordance with our pre-registration. We used a 2x3 repeated measures ANOVA to assess ratings of emotional salience for wins and losses with each partner. Additionally, in analyses involving trait differences between individuals, we utilized composite substance use scores and composite reward sensitivity scores, as well as squared reward sensitivity scores to further isolate aberrance associated with either extreme. In multiple linear regressions of behavioral data, we included differences in ratings for wins between each partner, as well as trait measures of substance use, reward sensitivity, aberrant reward sensitivity, and interaction terms between substance use and each of the two methods of assessing reward sensitivity. Although we initially anticipated the inclusion of IOS and its interaction with other terms in our multiple linear regression, due to the limited sample of data available for this measure, we chose to exclude it from the regression model.

Familywise error correction was applied only when multiple comparisons addressed a shared null hypothesis, such as in whole-brain voxelwise analyses (see below). Following recommendations from Rubin (2021), planned analyses were organized hierarchically— omnibus tests were conducted first, followed by simple effects or moderator analyses only when warranted. Because these tests evaluated distinct hypotheses, no additional correction for multiple comparisons was necessary.

### 2.6. Neuroimaging Data Collection

Functional images were acquired using a 3.0 Tesla Siemens PRISMA MRI scanner and a 20-channel head coil. Bold Oxygenation Level-Dependent (BOLD) sensitive functional images were acquired using a simultaneous multislice (multi-band factor = 2) gradient echo-planar imaging (EPI) sequence (240 mm in FOV, TR = 1,750 ms, TE = 29 ms, voxel size of 3.0 x 3.0 x 3.0 mm^3^, flip angle = 74°, interleaved slice acquisition, with 52 axial slices). Each run included 237 functional volumes. We also collected single-band reference images with each functional run of multi-band data to improve motion correction and registration. To facilitate anatomical localization and co-registration of functional data, a high-resolution structural scan was acquired (sagittal plane) with a T1-weighted magnetization-prepared rapid acquisition gradient echo (MPRAGE) sequence (224 mm in FOV, TR = 2,400 ms, TE = 2.17 ms, voxel size of 1.0 x 1.0 x 1.0 mm^3^, flip angle 8°). In addition, we also collected a B0 fieldmap to unwarp and undistort functional images (TR: 645 ms; TE1: 4.92 ms; TE2: 7.38 ms; matrix 74×74; voxel size: 2.97×2.97×2.80 mm; 58 slices, with 15% gap; flip angle: 60°).

### 2.7. Neuroimaging Preprocessing

Neuroimaging data were converted to the Brain Imaging Data Structure (BIDS) using HeuDiConv version (Halchenko et al., 2024). Results included in this manuscript come from preprocessing performed using fMRIPrep 20.2.3 (Esteban et al., 2019, 2018), which is based on Nipype 1.4.2 (K. Gorgolewski et al., 2011, 2018). The details described below are adapted from the fMRIPrep preprocessing details; extraneous details were omitted for clarity.

#### 2.7.1. Anatomical data preprocessing

The T1-weighted (T1w) image was corrected for intensity non-uniformity (INU) with ‘N4BiasFieldCorrection’,distributed with ANTs 2.3.3, and used as T1w-reference throughout the workflow. The T1w-reference was then skull-stripped with a Nipype implementation of the ‘antsBrainExtraction.sh’ workflow (from ANTs), using OASIS30ANTs as target template. Brain tissue segmentation of cerebrospinal fluid (CSF), white-matter (WM), and gray-matter (GM) was performed on the brain-extracted T1w using ‘fast’ (FSL 5.0.9). Volume-based spatial normalization to one standard space (MNI152NLin2009cAsym) was performed through nonlinear registration with ‘antsRegistration’ (ANTs 2.3.3), using brain-extracted versions of both T1w reference and the T1w template. The following template was selected for spatial normalization: ICBM 152 Nonlinear Asymmetrical template version 2009c (TemplateFlow ID: MNI152NLin2009cAsym)

#### 2.7.2. Functional data preprocessing

For each of the BOLD runs per participant, the following preprocessing steps were performed. First, a reference volume and its skull-stripped version were generated by aligning and averaging 1 single-band reference (SBRefs). A B0-nonuniformity map (or fieldmap) was estimated based on a phase-difference map calculated with a dual-echo GRE (gradient-recall echo) sequence, processed with a custom workflow of SDCFlows inspired by the ‘epidewarp.fsl’ script (http://www.nmr.mgh.harvard.edu/~greve/fbirn/b0/epidewarp.fsl) and further improvements in HCP Pipelines. The fieldmap then was co-registered to the target EPI (echo-planar imaging) reference run and converted to a displacements field map (amenable to registration tools such as ANTs) with FSL’s ‘fuguè and other SDCflows tools. Based on the estimated susceptibility distortion, a corrected EPI (echo-planar imaging) reference was calculated for a more accurate co-registration with the anatomical reference. The BOLD reference then was co-registered to the T1w reference using ‘flirt’ (FSL 5.0.9) with the boundary-based registration cost-function. Co-registration was configured with nine degrees of freedom to account for distortions remaining in the BOLD reference. Head-motion parameters with respect to the BOLD reference (transformation matrices, and six corresponding rotation and translation parameters) are estimated before any spatiotemporal filtering using ‘mcflirt’.

BOLD runs were slice-time corrected using ‘3dTshift’ from AFNI 20160207. First, a reference volume and its skull-stripped version were generated using a custom methodology of fMRIPrep. The BOLD time-series (including slice-timing correction when applied) were resampled onto their original, native space by applying a single, composite transform to correct for head-motion and susceptibility distortions. These resampled BOLD time-series will be referred to as preprocessed BOLD in original space, or just preprocessed BOLD. The BOLD time-series were resampled into standard space, generating a preprocessed BOLD run in MNI152NLin2009cAsym space. First, a reference volume and its skull-stripped version were generated using a custom methodology of fMRIPrep. Several confounding time-series were calculated based on the preprocessed BOLD, notably including framewise displacement (FD).

Additionally, a set of physiological regressors were extracted to allow for component-based noise correction (CompCor). Principal components are estimated after high-pass filtering the preprocessed BOLD time-series (using a discrete cosine filter with 128s cut-off) for anatomical component correction (aCompCor). For aCompCor, three probabilistic masks (CSF, WM and combined CSF+WM) were generated in anatomical space. The implementation differs from that of Behzadi et al. (2007) in that instead of eroding the masks by 2 pixels on BOLD space, the aCompCor masks are subtracted from a mask of pixels that likely contain a volume fraction of GM. This mask is obtained by thresholding the corresponding partial volume map at 0.05, and it ensures components are not extracted from voxels containing a minimal fraction of GM. Finally, these masks are resampled into BOLD space and binarized by thresholding at 0.99 (as in the original implementation). Components also are calculated separately within the WM and CSF masks. For each CompCor decomposition, the k components with the largest singular values are retained, such that the retained components’ time series are sufficient to explain 50 percent of variance across the nuisance mask (CSF, WM, combined, or temporal). The remaining components are dropped from consideration. The head-motion estimates calculated in the correction step also were placed within the corresponding confounds file. All resamplings can be performed with a single interpolation step by composing all the pertinent transformations (i.e., head-motion transform matrices, susceptibility distortion correction when available, and co-registrations to anatomical and output spaces). Gridded (volumetric) resamplings were performed using ‘antsApplyTransforms’ (ANTs), configured with Lanczos interpolation to minimize the smoothing effects of other kernels.

Many internal operations of fMRIPrep use Nilearn 0.6.2, mostly within the functional processing workflow. For more details of the pipeline, see the section corresponding to workflows in fMRIPrep’s documentation (https://fmriprep.readthedocs.io/en/latest/workflows.html).

Further, we applied spatial smoothing with a 5mm full-width at half-maximum (FWHM) Gaussian kernel using FMRI Expert Analysis Tool (FEAT) Version 6.00, part of FSL (FMRIB’s Software Library, www.fmrib.ox.ac.uk/fsl). Non-brain removal using BET (Smith, 2002) and grand mean intensity normalization of the entire 4D dataset by a single multiplicative factor were also applied.

### 2.8. Neuroimaging Analyses

All neuroimaging analyses were conducted in FSL version 6.0.4 (Jenkinson et al., 2012; Smith et al., 2004) using a general linear model (GLM) with local autocorrelation correction (Woolrich et al., 2001). We examined both task-evoked activation and task-modulated functional connectivity to test how BOLD responses were shaped by reward sensitivity and subclinical substance use. Following the modeling approach used in the Human Connectome Project (Barch et al., 2013), we treated each trial as a single event spanning the full decision and outcome phases. The activation model included ten regressors: win, loss, and neutral outcomes across the three partner conditions (friend, stranger, computer), plus one for missed trials. Each event was modeled as a 3.52-second boxcar beginning at trial onset.

Nuisance regressors included six motion parameters (translations and rotations), the first six aCompCor components explaining the most variance, framewise displacement, and non-steady state volumes. High-pass filtering was implemented using a discrete cosine transform with a 128-second cutoff. Participants completed two runs of the task (72 trials each), and contrast estimates were averaged across runs for group-level analyses. Full trial counts and event-level metadata are available in our open dataset and accompanying data descriptor (Smith et al., 2024).

Connectivity analyses were conducted using a generalized psychophysiological interaction (gPPI) framework (Friston et al., 1997; McLaren et al., 2012), with the bilateral ventral striatum (VS) from the Oxford-GSK-Imanova atlas (Tziortzi et al., 2011) as the seed. The VS time series was entered as a physiological regressor, and interaction terms were computed with each task condition to estimate context-dependent connectivity. This approach is supported by prior work demonstrating robust corticostriatal interactions during social and reward tasks (Di & Biswal, 2017; Smith et al., 2016; Smith & Delgado, 2017). Group-level PPI models used the same trait and nuisance regressors as the activation models.

Whole-brain group-level models included regressors for linear and quadratic reward sensitivity, substance use, their interaction, and nuisance covariates (mean framewise displacement and whole-brain tSNR from MRIQC; Esteban et al., 2017). Exploratory analyses were conducted using FSL’s Randomise tool (Winkler et al., 2014), with cluster correction applied at a voxelwise threshold of Z > 3.1 and familywise error–corrected significance at p < 0.05 (Worsley, 2001). Imaging results are visualized with MRIcroGL (Rorden, 2025).

To test for a replication of the partner × outcome interaction in ventral striatum, we conducted a 3 (partner: friend, stranger, computer) × 2 (outcome: win, loss) repeated-measures ANOVA on extracted β values from the bilateral ventral striatum ROI. This approach follows prior work (e.g., Fareri et al., 2012) and provides a direct test of within-subject effects across social contexts. Although this analysis also could be expressed using a regression model with effect-coded predictors, the underlying statistical model is identical; we retained the ANOVA format for consistency and interpretability. For planned contrasts (e.g., friend > stranger during reward), we computed difference scores and visualized pairwise effects. ROI-based PPI analyses focused on predefined regions implicated in social valuation and reward processing—posterior cingulate cortex (PCC), medial and ventromedial prefrontal cortex (mPFC, vmPFC), and posterior temporoparietal junction (pTPJ)—defined using the Oxford-GSK-Imanova atlas and prior literature (Bhanji et al., 2019; Mars et al., 2012b).

### 2.9. Deviations from Pre-Registration

As noted in the Introduction and Participants section, several deviations from our preregistered procedures occurred, due in part to constraints imposed by the COVID-19 pandemic. We initially planned to recruit 100 college-aged participants (18–22 years) but ultimately collected usable data from 45 participants. We also intended to collect both emotional salience ratings and perceived closeness ratings (Inclusion of Other in Self; IOS; Aron et al., 1992) for each partner (friend, stranger, computer) during the experimental session. Whereas emotional salience ratings were collected as planned, IOS ratings for the stranger and computer conditions were inadvertently omitted during scanning and instead collected during a delayed follow-up with 28 participants. Given the variability in follow-up timing, the reduced sample size, and the potential for memory-related bias, we excluded IOS scores from all analyses.

Although our preregistration included exploratory analyses examining self-reported social closeness as a moderator of neural responses to shared rewards, we were unable to conduct these analyses due to the incomplete IOS data collection described above.

We also preregistered hypotheses examining interactions between substance use and closeness ratings (IOS) in order to assess effects independent of reward sensitivity. However, the sample showed a limited range of substance use, restricting our ability to evaluate these hypotheses meaningfully. Although we retained our composite substance use score as a covariate in relevant models, our results focus more heavily on the preregistered whole-brain activation analysis and an exploratory ROI connectivity analysis, rather than on substance use moderation effects.

Additionally, although our preregistration identified reward sensitivity and substance use as key moderators, it did not specify how these constructs would be operationalized. As detailed in the Individual Differences Measures section, we collected two self-report instruments for each trait and created composite indices by z-scoring and summing the measures, followed by decile binning to reduce the influence of extreme values. These composites were entered into the models as planned, and we also report their interaction, which was not preregistered but emerged as theoretically meaningful in interpreting individual differences.

## 3. Results

### 3.1. Reward Sensitivity Moderates Preference for Sharing Wins with Friends

We first examined whether individual differences in reward sensitivity and substance use predicted subjective emotional responses to shared rewards across different social contexts. Emotional salience ratings showed a significant Partner × Outcome interaction (F(2, 78) = 35.277, p < 0.001, ηp² = 0.475), indicating that the effect of social partner on emotional ratings differed between win and loss outcomes. Post-hoc comparisons (Bonferroni-corrected) confirmed that participants rated winning with a friend as significantly more positive (M = 3.65, SD = 1.83) compared to winning with a stranger (M = 0.77, SD = 2.42; t(39) = -7.205, p < .001, d = -1.298) or a computer (M = 0.53, SD = 2.76; t(39) = -6.382, p < .001, d = -1.261). Likewise, participants rated losing with a friend as significantly more negative (M = -2.86, SD = 2.38) compared to losses with a stranger (M = -1.56, SD = 1.85; t(39) = 3.261, p = .007, d = 0.627) or computer (M = -1.47, SD = 2.06; t(39) = 3.872, p = .001, d = 0.673). Ratings for stranger and computer partners did not differ significantly in either outcome condition. These findings confirm that the identity of the social interaction partner significantly shaped participants’ emotional experiences.

Next, we tested whether these differences in emotional salience across social contexts were moderated by individual differences in reward sensitivity and substance use. Specifically, we examined a nonlinear regression model predicting differences in emotional salience ratings between winning with a close friend versus a computer, including reward sensitivity (linear and quadratic terms), substance use, and their interactions. This model revealed a significant quadratic effect of reward sensitivity (t(41) = 2.381, p = 0.046; Fig. 2), indicating that both relatively high and low reward sensitivity scores were associated with a stronger preference for winning with friends relative to computers. A similar quadratic effect was observed when examining win–loss differentials for friends versus computers (t(41) = 2.341, p = 0.025). Notably, no significant moderation effects were found for the friend versus stranger contrasts, suggesting that this quadratic relationship may be specific to comparisons between friends and non-social partners. These findings indicate that individuals with extreme levels of reward sensitivity assign greater interpersonal salience to rewards when shared with close social partners compared to non-social contexts.

**Figure 2.**
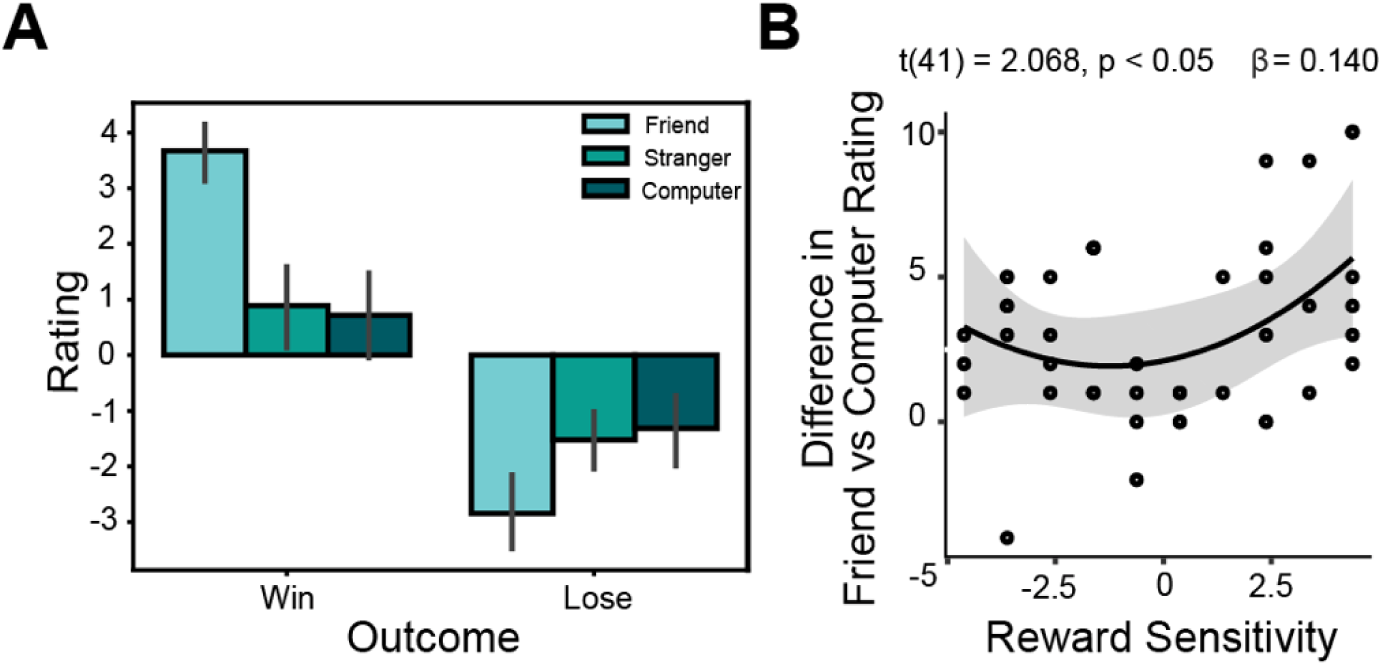
Aberrant reward sensitivity is associated with strength of preference for winning with friends. (A) Participants rated the emotional salience of winning and losing with each partner. An ANOVA demonstrated that ratings for winning and losing were significantly different across partners; the highest mean emotional salience in both conditions was with friends. (B) Participants in our sample who were either hyposensitive or hypersensitive to rewards (aberrant reward sensitivity) had a significantly greater boost in how they rated winning with a close friend, as opposed to a non-human partner, than those with more average reward sensitivity. However, reward sensitivity was not significantly associated with a difference in ratings of wins with friends as opposed to human strangers.

### 3.2. Reward Sensitivity Moderates Enhancement of VS Activation in Response to Close Friends

To examine whether reward sensitivity and substance use moderated neural responses to shared rewards, we analyzed ventral striatum (VS) activation during the outcome phase of the task. Based on prior findings (Fareri et al., 2012), we compared reward-related VS activation across the friend versus stranger and friend versus computer conditions. A repeated-measures ANOVA revealed a significant interaction between social partner and outcome type (F(1.56, 68.86) = 3.676, p = 0.041, ηp² = .077, representing a medium effect size; Fig. 3B). Post-hoc comparisons (Bonferroni-corrected across three partner contrasts) showed greater VS activation when participants shared rewards with friends compared to computers (t=2.935, p=0.0055) and compared to strangers (t=2.210, p=0.0422). No significant differences were observed for loss outcomes, indicating that partner effects were specific to rewarding outcomes.

**Figure 3.**
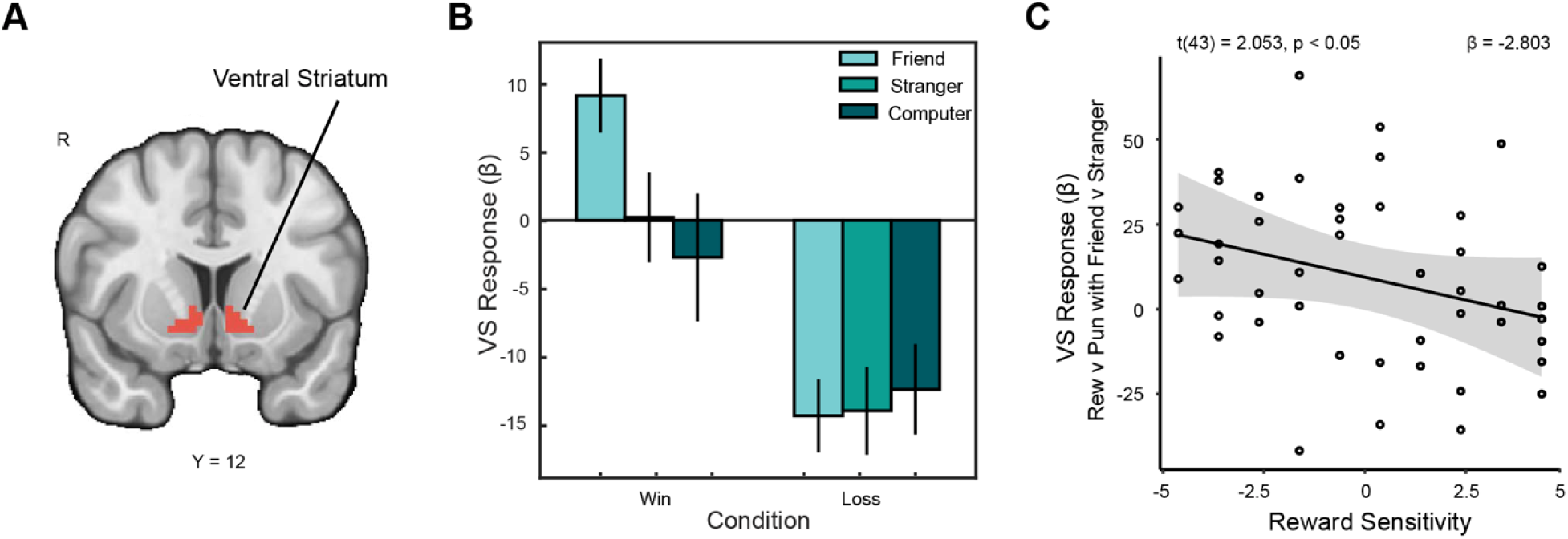
Ventral striatal response to reward with close friends is associated with reward sensitivity. (A) We focused on a pre-registered ventral striatal (VS) region of interest (Tziortzi et al., 2011). (B) Within this region of interest, VS activation was significantly greater during wins with friends than wins with strangers. (C) We conducted a pre-registered analysis to determine whether reward sensitivity independently moderated VS activation for reward as opposed to punishment with friends as opposed to strangers. We found that as trait reward sensitivity increases, the overall difference in VS activation for outcomes shared with friends relative to strangers decreases. Those in our sample with low trait reward sensitivity showed a greater difference in VS enhancement when the context was a closer social relation than those with high reward sensitivity.

We next tested whether this partner-related VS activation difference was moderated by individual differences in reward sensitivity and substance use. Regression models included linear and quadratic terms for reward sensitivity to test for potential nonlinear effects, as well as substance use and their interaction. Within this model, a significant moderation by reward sensitivity was observed (β = -2.80, SE = 1.37, t = -2.053, p = 0.047, f^2^ = 0.114, representing a small effect size; Fig. 3C). Specifically, individuals higher in reward sensitivity exhibited less differentiation in VS activation between rewards shared with friends versus strangers, whereas individuals lower in reward sensitivity showed stronger differentiation. No moderation effects were observed for substance use (t = -1.009, p = .319) or quadratic reward sensitivity (t = - 0.635, p = .529) in this analysis. These results suggest that trait-level reward sensitivity modulates how selectively the VS encodes reward outcomes in socially meaningful contexts.

To complement the ROI analyses, we conducted a whole-brain analysis contrasting friend > stranger conditions during reward outcomes, revealing a significant cluster extending into the VS (x = 8.5, y = -0.5, z = -3.5; ke = 53), which further supports the VS’s involvement in social reward processing. Additionally, we examined whether substance use moderated neural responses to social reward elsewhere in the brain. A significant interaction emerged in the superior temporal sulcus (STS), such that higher sub-clinical substance use predicted greater STS activation for the friend > computer reward contrast (x = 54, y = -40, z = 11; ke = 37; t(43) = 3.783, p = 0.0005). These results indicate heightened neural sensitivity to shared social rewards among individuals with higher levels of substance use, particularly in regions linked to social cognition.

### 3.3. Sharing Rewards with Close Friends Enhances Corticostriatal Connectivity

To test whether social partners influenced functional connectivity with the ventral striatum, we conducted seed-to-target psychophysiological interaction (PPI) analyses using the VS as a seed region and the posterior TPJ (pTPJ) as a target region. A 2 (outcome) x 3 (partner) repeated-measures ANOVA on VS–pTPJ connectivity revealed a significant main effect of social partner (F(2, 88) = 7.841, p = 0.0007, ηp²=0.151, representing a large effect size; Fig. 4). Follow-up tests showed significantly greater VS–pTPJ connectivity for friend > stranger (t(44) = 2.59, p = 0.038) and friend > computer (t(44) = -3.02, p = 0.013) contrasts during reward outcomes. We emphasized the friend > stranger contrast for TPJ analyses because perceived social closeness and human intentionality are best captured through comparisons between two human agents. This aligns with prior work highlighting the TPJ’s role in mentalizing and social cognition (Saxe & Kanwisher, 2013). The computer condition, though nonsocial, lacks the interpersonal variability that defines theory-of-mind processing. Overall, these results indicate that positive, socially meaningful experiences enhance corticostriatal connectivity during reward receipt. No significant task-modulated connectivity emerged between the VS and other pre-specified ROIs (vmPFC, mPFC, PCC).

**Figure 4.**
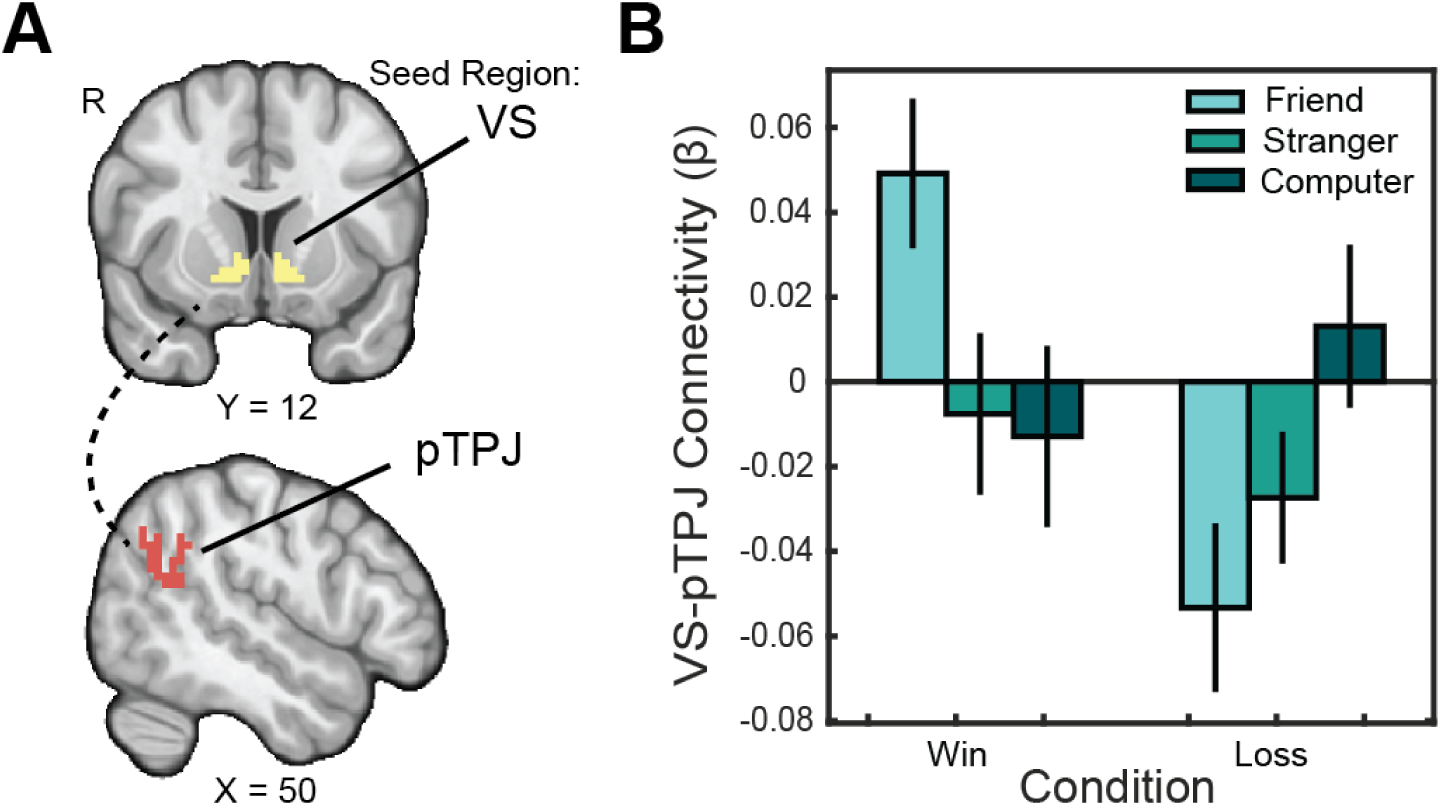
Social closeness enhances connectivity between VS-TPJ. (A) We pre-registered Regions of Interest (ROIs) in the VS (Tziortzi et al., 2011) and the posterior TPJ (Mars et al., 2012b). (B) We ran an analysis examining psychophysiological interactions between the VS seed and pTPJ target region to further probe for group differences in the effects of partner on reward processing. Connectivity between these regions for reward as opposed to punishment was significantly enhanced in the friend condition as opposed to the stranger condition.

To assess the role of individual differences in moderating VS-pTPJ connectivity, we regressed VS–pTPJ connectivity on substance use, reward sensitivity, and their interaction. Based on prior findings linking substance use to alterations in social valuation (e.g., Franken & Muris, 2006; Kober et al., 2010), we hypothesized that higher levels of substance use would enhance VS– pTPJ connectivity during socially rewarding contexts involving close friends. Consistent with our hypothesis, sub-clinical substance use was a significant predictor of greater VS–pTPJ connectivity for the friend > stranger reward contrast (β = 0.068, SE = 0.027, t = 2.482, p = 0.017, f^2^ = 0.171, representing a medium effect size), independent of reward sensitivity.

To extend our ROI findings, we conducted exploratory whole-brain PPI analyses to examine whether VS connectivity with other regions varied as a function of social partner and individual differences. We prioritized the friend > computer contrast for analyses involving perceptual salience and facial processing regions such as the fusiform face area (FFA). This allowed us to isolate the effect of a socially meaningful human partner relative to a perceptually similar but nonsocial control. These analyses revealed enhanced connectivity between the VS and the right fusiform face area (rFFA; x = 44.5, y = –45.5, z = –21.5; ke = 29; Fig. 5) and bilateral occipital cortex (x = –39.5, y = –81.5, z = –12.5; ke = 41) when participants shared rewards with a close friend compared to a computer partner. Notably, reward sensitivity moderated these effects (rFFA: β = 0.034, SE = 0.012, t = 2.845, p = 0.007, f² = 0.219; occipital: β = 0.018, SE = 0.007, t = 2.670, p = 0.011, f^2^ = 0.193), with higher reward sensitivity associated with stronger connectivity. Substance use also independently moderated VS–rFFA connectivity (β = 0.057, SE = 0.026, t(43) = 2.213, p = 0.033, f² = 0.132). Together, these exploratory findings suggest that individual differences in reward sensitivity and substance use may jointly shape how reward signals interact with perceptual and social-processing regions during close social interactions, consistent with evidence that VS–FFA coupling supports motivated responses to socially salient stimuli (Genevsky et al., 2017).

**Figure 5.**
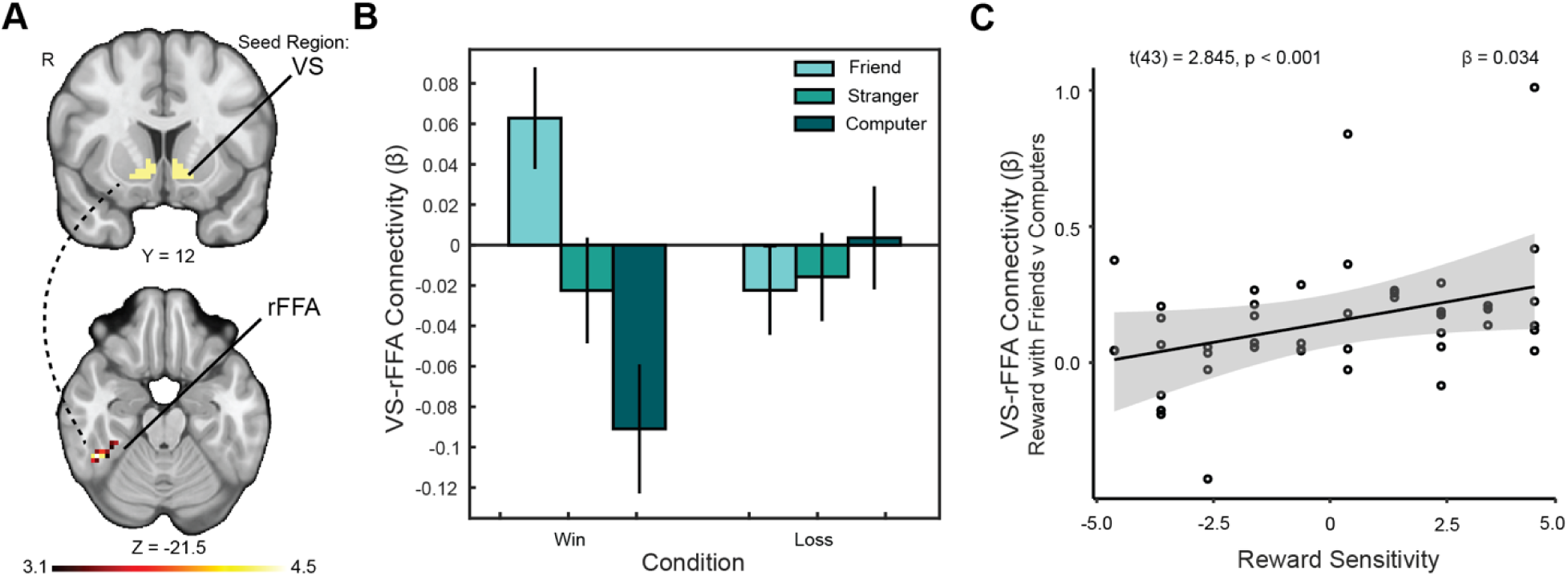
VS-rFFA connectivity is moderated by reward sensitivity. (A) We identified a cluster extending into the right temporo-occipital fusiform face area (rFFA) for which connectivity with the ventral striatum (VS) was significantly greater when sharing rewards with friends as opposed to computers, reflecting increased engagement of perceptual-social circuits in response to close social partners. The overlaying Z statistic image was generated using cluster-forming thresholding of Z > 3.1 with a family-wise error rate of P = 0.05. (B) Connectivity between the VS and rFFA increased significantly when participants viewed a close friend’s face during reward receipt, relative to a perceptually similar but nonsocial partner. (C) Upon discovery of the rFFA cluster, we conducted an exploratory analysis to test whether this connectivity was moderated by reward sensitivity, independent of substance use. We found that the enhancement was significantly moderated by trait reward sensitivity: participants with higher reward sensitivity showed a greater increase in VS–rFFA connectivity when sharing rewards with friends relative to computers.

## 4. Discussion

The present study examined how social context modulates neural responses to shared rewards as a function of individual differences in trait reward sensitivity and sub-clinical substance use. Our results replicate prior findings that close social partners enhance ventral striatal (VS) activation during reward receipt (Fareri et al., 2012) and extend this work by showing that individual traits moderate these effects. Specifically, reward sensitivity predicted the extent to which neural activation and corticostriatal connectivity differed between close friends and other partners. Sub-clinical substance use, though uncorrelated with reward sensitivity, also moderated neural responses, particularly in regions implicated in social cognition. These findings suggest that social context, personality traits, and substance use history jointly shape neural responses to reward, even in non-clinical populations.

Our results align with prior work linking striatal activity to both primary and social rewards (Knutson et al., 2000; O’Doherty et al., 2017), and with research identifying the VS as sensitive to social context (Fareri et al., 2022; Lockwood et al., 2020). We found that individuals lower in reward sensitivity exhibited stronger VS activation when winning with friends relative to strangers, consistent with a selective sensitivity to close relationships. In contrast, individuals with higher reward sensitivity showed more uniform responses across human partners, suggesting that they may be broadly responsive to social reward. Importantly, reward sensitivity also moderated corticostriatal connectivity with the fusiform face area (FFA) and the posterior temporoparietal junction (pTPJ)—regions implicated in face processing and theory of mind, respectively (Deen et al., 2015; Park et al., 2017). These findings suggest that heightened trait reward sensitivity may enhance social valuation mechanisms, particularly in visually salient and personally relevant contexts.

Notably, the effects observed for shared reward did not generalize to loss outcomes. This asymmetry is consistent with evidence that the ventral striatum is more strongly activated by appetitive outcomes and gain-related signals (Knutson et al., 2001; Delgado, 2007), particularly in social contexts involving cooperation or mutual benefit (Fareri & Delgado, 2014). In contrast, aversive or negative outcomes such as monetary losses or outcomes in competitive social contexts (Delgado et al., 2008) often are associated with activation in regions like the anterior insula and dorsal anterior cingulate cortex (dACC), which support salience detection, emotional conflict, and affective pain processing (Eisenberger, 2012; Wager et al., 2016). These regions may be less modulated by interpersonal closeness, helping explain why trait- and context-based differences were most evident during positive outcomes. Alternatively, from a computational perspective, the relative difference between reward and punishment signals may be more behaviorally relevant than absolute responses to either outcome type alone, suggesting that social modulation effects may be most pronounced when examining reward-punishment contrasts rather than isolated responses.

We also observed that sub-clinical substance use levels independently moderated neural responses to reward. Substance use was linked with increased corticostriatal connectivity with both the pTPJ and FFA when sharing rewards with friends. These regions have been previously implicated in both social cognition (Mars et al., 2012a; Saxe & Kanwisher, 2013; Young et al., 2007) and cue responsivity in substance use contexts (Groefsema et al., 2020). These findings suggest that even in non-clinical populations, individual differences in motivational traits and substance use behaviors may jointly influence neural circuits engaged during rewarding social interactions. Although correlational, these results are consistent with prior behavioral work suggesting that peer presence and social salience enhance the appeal of rewarding stimuli (Shadur & Hussong, 2014; Volkow et al., 2011).

Several limitations warrant consideration. Due to experimenter error, we were underpowered to assess pre-registered analyses involving subjective closeness ratings and could not examine whether perceived closeness moderated our neural findings. Although we included a broad range of reward sensitivity and substance use scores, our sample was restricted to healthy young adults and excluded individuals with clinically significant substance use. Further, although this design choice may limit generalizability to clinical populations, our goal was to examine dimensional variability in motivational traits within a non-clinical range. We believe these effects are valuable for identifying potential neural mechanisms that precede clinical impairment, consistent with dimensional approaches to risk (e.g., RDoC framework). Future work should assess whether similar patterns are observed in adolescent or older adult populations, where age-related changes in social motivation and striatal reactivity are well documented (Gadassi Polack et al., 2023; Telzer et al., 2013). Moreover, longitudinal designs will be needed to clarify the directional and predictive value of these neural markers for emerging psychopathology. In addition, although our modeling approach follows that used in large-scale datasets such as the Human Connectome Project gambling task (Barch et al., 2013), the short, fixed ITI necessarily limits the ability to isolate responses specific to reward consumption. Thus, our interpretations focus on relative differences across social contexts rather than on temporally discrete psychological processes.

Finally, given the modest sample size and analytic deviations from our preregistration, future studies will be important for validating these findings. We hope our open data and materials provide a useful foundation for that work (Smith et al., 2024). We also note that although different scoring strategies could be applied to the reward sensitivity and substance use measures—for example, using a single scale, raw z-scores, or alternative binning approaches— the high correlations between scales suggest that such choices would not meaningfully affect the pattern of results. Our analytic approach prioritized parsimony and alignment with theoretical goals (e.g., modeling both linear and quadratic effects), and we believe the results are robust to reasonable variations in scaling. To promote transparency and reproducibility, all de-identified data and analysis code are openly available via GitHub and OpenNeuro (Markiewicz et al., 2021), allowing others to explore alternative operationalizations and extensions of the current work.

Despite these limitations, the present findings contribute to a growing literature on how trait-level characteristics shape neural responses to social rewards. Our results demonstrate that close social relationships selectively engage corticostriatal circuits, and that these effects are amplified or altered by individual differences in reward sensitivity and sub-clinical substance use. Given prior associations between aberrant reward sensitivity, social cognition, and mental health risk (Alloy et al., 2016; Nusslock et al., 2012), these findings point to potential neurobiological pathways through which social context and individual traits confer vulnerability or resilience. Social contexts are known to amplify ventral striatal responses and approach motivation, particularly in the presence of peers (Chein et al., 2011; Telzer et al., 2015), and heightened sensitivity to socially rewarding cues has been linked to elevated substance-use risk (Beard et al., 2022; Sazhin et al., 2020). Recent work further suggests that dopamine system function modulates the association between ventral striatal responses to social reward and substance-use severity, such that diminished dopaminergic tone strengthens the link between blunted social reward responsivity and greater substance use (Jarcho et al., 2022). Taken together, these considerations suggest that individual differences in social reward processing may help explain why social isolation, peer influence, and substance use often intersect in young adulthood, and they highlight the potential value of interventions that strengthen social connectedness or modulate reward motivation in at-risk populations.

## Acknowledgments

This work was supported in part by grants from the National Institute of Mental Health (R01-MH123473 and R01-MH126911 to LBA, R15-MH122927 to DSF), the Eunice Kennedy Shriver National Institute of Child Health and Human Development (R21-HD093912 to JMJ), the National Institute on Aging (R01-AG067011 to DVS and F31-AG085934 to JBW), and the National Institute on Drug Abuse (R03-DA046733 to DVS), and a fellowship from the Temple Public Policy Lab (to JMJ).

## Data and code availability

Analysis code related to this project can be found on GitHub (https://github.com/DVS-Lab/istart-sharedreward). In addition, all statistical maps (thresholded and unthresholded) can be found on NeuroVault (https://identifiers.org/neurovault.collection:15006); and raw data can be found on OpenNeuro (https://openneuro.org/datasets/ds004920/versions/1.1.1).

## References

Acikalin, M. Y., Gorgolewski, K. J., & Poldrack, R. A. (2017). A Coordinate-Based Meta-Analysis of Overlaps in Regional Specialization and Functional Connectivity across Subjective Value and Default Mode Networks. Frontiers in Neuroscience, 11(1). doi:10.3389/fnins.2017.00001

Ait Oumeziane, B., Jones, O., & Foti, D. (2019). Neural Sensitivity to Social and Monetary Reward in Depression: Clarifying General and Domain-Specific Deficits. Frontiers in Behavioral Neuroscience, 13. https://www.frontiersin.org/articles/10.3389/fnbeh.2019.00199

Alloy, L. B., Abramson, L. Y., Walshaw, P. D., Cogswell, A., Smith, J. M., Neeren, A. M., Hughes, M. E., Iacoviello, B. M., Gerstein, R. K., Keyser, J., Urosevic, S., & Nusslock, R. (2006). Behavioral Approach System (BAS) Sensitivity and Bipolar Spectrum Disorders: A Retrospective and Concurrent Behavioral High-Risk Design. Motivation and Emotion, 30(2), 143–155. 10.1007/s11031-006-9003-3

Alloy, L. B., Bender, R. E., Wagner, C. A., Whitehouse, W. G., Abramson, L. Y., Hogan, M. E., Sylvia, L. G., & Harmon-Jones, E. (2009). Bipolar Spectrum – Substance Use Co-occurrence: Behavioral Approach System (BAS) Sensitivity and Impulsiveness as Shared Personality Vulnerabilities. Journal of Personality and Social Psychology, 97(3), 549–565. 10.1037/a0016061

Alloy, L. B., Bender, R. E., Whitehouse, W. G., Wagner, C. A., Liu, R. T., Grant, D. A., Jager-Hyman, S., Molz, A., Choi, J. Y., Harmon-Jones, E., & Abramson, L. Y. (2012). High Behavioral Approach System (BAS) Sensitivity, Reward Responsiveness, and Goal-Striving Predict First Onset of Bipolar Spectrum Disorders: A Prospective Behavioral High-Risk Design. Journal of Abnormal Psychology, 121(2), 339–351. 10.1037/a0025877

Alloy, L. B., Olino, T., Freed, R. D., & Nusslock, R. (2016). Role of Reward Sensitivity and Processing in Major Depressive and Bipolar Spectrum Disorders. Behavior Therapy, 47(5), 600–621. 10.1016/j.beth.2016.02.014

Aron, A., Aron, E. N., & Smollan, D. (1992). Inclusion of Other in the Self Scale and the structure of interpersonal closeness. Journal of Personality and Social Psychology, 63(4), 596–612. 10.1037/0022-3514.63.4.596

Barch, D. M., Burgess, G. C., Harms, M. P., Petersen, S. E., Schlaggar, B. L., Corbetta, M., Glasser, M. F., Curtiss, S., Dixit, S., Feldt, C., Nolan, D., Bryant, E., Hartley, T., Footer, O., Bjork, J. M., Poldrack, R., Smith, S., Johansen-Berg, H., Snyder, A. Z., & Van Essen, D. C. (2013). Function in the human connectome: Task-fMRI and individual differences in behavior. NeuroImage, 80, 169–189. 10.1016/j.neuroimage.2013.05.033

Bart, C. P., Titone, M. K., Ng, T. H., Nusslock, R., & Alloy, L. B. (2021). Neural reward circuit dysfunction as a risk factor for bipolar spectrum disorders and substance use disorders: A review and integration. Clinical Psychology Review, 87, 102035. 10.1016/j.cpr.2021.102035

Bartra, O., McGuire, J. T., & Kable, J. W. (2013). The valuation system: a coordinate-based meta-analysis of BOLD fMRI experiments examining neural correlates of subjective value. NeuroImage, 76, 412–427. doi:10.1016/j.neuroimage.2013.02.063

Basedow, L. A., Kuitunen-Paul, S., Eichler, A., Roessner, V., & Golub, Y. (2021). Diagnostic Accuracy of the Drug Use Disorder Identification Test and Its Short Form, the DUDIT-C, in German Adolescent Psychiatric Patients. Frontiers in Psychology, 12, 678819. 10.3389/fpsyg.2021.678819

Beard, S. J., Yoon, L., Venticinque, J. S., Shepherd, N. E., & Guyer, A. E. (2022). The brain in social context: A systematic review of substance use and social processing from adolescence to young adulthood. Developmental Cognitive Neuroscience, 57, 101147. 10.1016/j.dcn.2022.101147

Behzadi, Y., Restom, K., Liau, J., & Liu, T. T. (2007). A component based noise correction method (CompCor) for BOLD and perfusion based fMRI. NeuroImage, 37(1), 90–101.

Berman, A. H., Bergman, H., Palmstierna, T., & Schlyter, F. (n.d.). Drug Use Disorders Identification Test (DUDIT). APA PsycTests. 10.1037/t02890-000

Berridge, K. C., & Robinson, T. E. (2016). Liking, wanting, and the incentive-sensitization theory of addiction. American Psychologist, 71(8), 670–679. 10.1037/amp0000059

Bhanji, J. P., & Delgado, M. R. (2014). The social brain and reward: Social information processing in the human striatum. WIREs Cognitive Science, 5(1), 61–73. 10.1002/wcs.1266

Bhanji, J., Smith, D. V., & Delgado, M. (2019). A Brief Anatomical Sketch of Human Ventromedial Prefrontal Cortex [Preprint]. PsyArXiv. 10.31234/osf.io/zdt7f

Bolis, D., Lahnakoski, J. M., Seidel, D., Tamm, J., & Schilbach, L. (2021). Interpersonal similarity of autistic traits predicts friendship quality. Social Cognitive and Affective Neuroscience, 16(1–2), 222–231. 10.1093/scan/nsaa147

Brooks, S. J., Dalvie, S., Cuzen, N. L., Cardenas, V., Fein, G., & Stein, D. J. (2014). Childhood adversity is linked to differential brain volumes in adolescents with alcohol use disorder: A voxel-based morphometry study. Metabolic Brain Disease, 29(2), 311–321. 10.1007/s11011-014-9489-4

Büchel, C., Holmes, A. P., Rees, G., & Friston, K. J. (1998). Characterizing stimulus-response functions using nonlinear regressors in parametric fMRI experiments. NeuroImage, 8(2), 140–148. 10.1006/nimg.1998.0351

Büchel, C., Peters, J., Banaschewski, T., Bokde, A. L. W., Bromberg, U., Conrod, P. J., Flor, H., Papadopoulos, D., Garavan, H., Gowland, P., Heinz, A., Walter, H., Ittermann, B., Mann, K., Martinot, J.-L., Paillère-Martinot, M.-L., Nees, F., Paus, T., Pausova, Z., … IMAGEN consortium. (2017). Blunted ventral striatal responses to anticipated rewards foreshadow problematic drug use in novelty-seeking adolescents. Nature Communications, 8, 14140. 10.1038/ncomms14140

Castrellon, J. J., Young, J. S., Dang, L. C., Cowan, R. L., Zald, D. H., & Samanez-Larkin, G. R. (2019). Mesolimbic dopamine D2 receptors and neural representations of subjective value. Scientific Reports, 9(1), 20229. doi:10.1038/s41598-019-56858-1

Carver, C. S., & White, T. L. (1994). Behavioral inhibition, behavioral activation, and affective responses to impending reward and punishment: The BIS/BAS Scales. Journal of Personality and Social Psychology, 67(2), 319–333. 10.1037/0022-3514.67.2.319

Chat, I. K.-Y., Dunning, E. E., Bart, C. P., Carroll, A. L., Grehl, M. M., Damme, K. S. F., Abramson, L. Y., Nusslock, R., & Alloy, L. B. (2022). The Interplay Between Reward-Relevant Life Events and Trait Reward Sensitivity in Neural Responses to Reward Cues. Clinical Psychological Science, 10(5), 869–884. 10.1177/21677026211056627

Chein, J., Albert, D., O’Brien, L., Uckert, K., & Steinberg, L. (2011). Peers increase adolescent risk taking by enhancing activity in the brain’s reward circuitry. Dev Sci, 14(2), F1–10. doi:10.1111/j.1467-7687.2010.01035.x

Deen, B., Koldewyn, K., Kanwisher, N., & Saxe, R. (2015). Functional organization of social perception and cognition in the superior temporal sulcus. Cerebral Cortex, 25(11). 10.1093/cercor/bhv111

Delgado, M. R., Nystrom, L. E., Fissell, C., Noll, D. C., & Fiez, J. A. (2000). Tracking the Hemodynamic Responses to Reward and Punishment in the Striatum. Journal of Neurophysiology, 84(6), 3072–3077. 10.1152/jn.2000.84.6.3072

Dennison, J. B., Sazhin, D., & Smith, D. V. (2022). Decision neuroscience and neuroeconomics: Recent progress and ongoing challenges. WIREs Cognitive Science, 13(3). 10.1002/wcs.1589

Di, X., & Biswal, B. B. (2017). Psychophysiological Interactions in a Visual Checkerboard Task: Reproducibility, Reliability, and the Effects of Deconvolution. Frontiers in Neuroscience, 11, 573. 10.3389/fnins.2017.00573

Dobryakova, E., & Smith, D. V. (2022). Reward enhances connectivity between the ventral striatum and the default mode network. NeuroImage, 258, 119398. 10.1016/j.neuroimage.2022.119398

Doricchi, F., Lasaponara, S., Pazzaglia, M., & Silvetti, M. (2022). Left and right temporal-parietal junctions (TPJs) as “match/mismatch” hedonic machines: A unifying account of TPJ function. Physics of Life Reviews, 42, 56–92. 10.1016/j.plrev.2022.07.001

Dziura, S. L., McNaughton, K. A., Giacobbe, E., Yarger, H., Hickey, A., Shariq, D., & Redcay, E. (2022). Neural sensitivity to social reward predicts social behavior and satisfaction in adolescents during the COVID-19 pandemic [Preprint]. PsyArXiv. 10.31234/osf.io/y9t8g

Esteban, O., Birman, D., Schaer, M., Koyejo, O. O., Poldrack, R. A., & Gorgolewski, K. J. (2017). MRIQC: Advancing the automatic prediction of image quality in MRI from unseen sites. PLOS ONE, 12(9), e0184661. 10.1371/journal.pone.0184661

Esteban, O., Markiewicz, C. J., Blair, R. W., Moodie, C. A., Isik, A. I., Erramuzpe, A., Kent, J. D., Goncalves, M., DuPre, E., Snyder, M., Oya, H., Ghosh, S. S., Wright, J., Durnez, J., Poldrack, R. A., & Gorgolewski, K. J. (2019). fMRIPrep: A robust preprocessing pipeline for functional MRI. Nature Methods, 16(1), 111–116. 10.1038/s41592-018-0235-4

Esteban, Oscar, Markiewicz, Christopher J., Goncalves, Mathias, Provins, Céline, Kent, James D., DuPre, Elizabeth, Salo, Taylor, Ciric, Rastko, Pinsard, Basile, Blair, Ross W., Poldrack, Russell A., & Gorgolewski, Krzysztof J. (2018). fMRIPrep: A robust preprocessing pipeline for functional MRI (23.1.3) [Computer software]. Zenodo. 10.5281/ZENODO.852659

Fareri, D. S., Chang, L. J., & Delgado, M. R. (2015). Computational Substrates of Social Value in Interpersonal Collaboration. The Journal of Neuroscience, 35(21), 8170–8180. 10.1523/JNEUROSCI.4775-14.2015

Fareri, D. S., & Delgado, M. R. (2014). Social Rewards and Social Networks in the Human Brain. The Neuroscientist, 20(4), 387–402. 10.1177/1073858414521869

Fareri, D. S., Hackett, K., Tepfer, L. J., Kelly, V., Henninger, N., Reeck, C., Giovannetti, T., & Smith, D. V. (2022). Age-related differences in ventral striatal and default mode network function during reciprocated trust. NeuroImage, 256. 10.1016/j.neuroimage.2022.119267

Fareri, D. S., Niznikiewicz, M. A., Lee, V. K., & Delgado, M. R. (2012). Social network modulation of reward-related signals. Journal of Neuroscience, 32(26). 10.1523/JNEUROSCI.0610-12.2012

Franken, I. H. A., & Muris, P. (2006). BIS/BAS personality characteristics and college students’ substance use. Personality and Individual Differences, 40(7). 10.1016/j.paid.2005.12.005

Friston, K. J., Buechel, C., Fink, G. R., Morris, J., Rolls, E., & Dolan, R. J. (1997). Psychophysiological and Modulatory Interactions in Neuroimaging. NeuroImage, 6(3), 218–229. 10.1006/nimg.1997.0291

Gadassi Polack, R., Mollick, J. A., Keren, H., Joormann, J., & Watts, R. (2023). Neural responses to reward valence and magnitude from pre- to early adolescence. NeuroImage, 275, 120166. 10.1016/j.neuroimage.2023.120166

Genevsky, A., Yoon, C., & Knutson, B. (2017). When brain beats behavior: Neuroforecasting crowdfunding outcomes. Journal of Neuroscience, 37(36). 10.1523/JNEUROSCI.1633-16.2017

Gorgolewski, K., Burns, C., Madison, C., Clark, D., Halchenko, Y., Waskom, M., & Ghosh, S. (2011). Nipype: A Flexible, Lightweight and Extensible Neuroimaging Data Processing Framework in Python. Frontiers in Neuroinformatics, 5. https://www.frontiersin.org/articles/10.3389/fninf.2011.00013

Gorgolewski, K. J., Esteban, O., Ellis, D. G., Notter, M. P., Ziegler, E., Johnson, H., Hamalainen, C., Yvernault, B., Burns, C., Manhães-Savio, A., Jarecka, D., Markiewicz, C. J., Salo, T., Clark, D., Waskom, M., Wong, J., Modat, M., Dewey, B. E., Clark, M. G., … Ghosh, S. (2018). Nipype: A flexible, lightweight and extensible neuroimaging data processing framework in Python. 0.13.1 [Computer software]. Zenodo. https://zenodo.org/record/581704

Gotlib, I. H., Hamilton, J. P., Cooney, R. E., Singh, M. K., Henry, M. L., & Joormann, J. (2010). Neural processing of reward and loss in girls at risk for major depression. Archives of General Psychiatry, 67(4), 380–387. 10.1001/archgenpsychiatry.2010.13

Groefsema, M. M., Mies, G. W., Cousijn, J., Engels, R. C. M. E., Sescousse, G., & Luijten, M. (2020). Brain responses and approach bias to social alcohol cues and their association with drinking in a social setting in young adult males. The European Journal of Neuroscience, 51(6), 1491–1503. 10.1111/ejn.14574

Haber, S. N., & Knutson, B. (2010). The Reward Circuit: Linking Primate Anatomy and Human Imaging. Neuropsychopharmacology, 35(1), 4–26. 10.1038/npp.2009.129

Haeger, A., Lee, H., Fell, J., & Axmacher, N. (2014). Selective processing of buildings and faces during working memory: The role of the ventral striatum. European Journal of Neuroscience, 41(4), 505–513. 10.1111/ejn.12808

Halchenko, Y. O., Goncalves, M., Ghosh, S., Velasco, P., Castello, M., Salo, T., Wodder, J., Hanke, M., Sadil, P., Gorgolewski, K., Ioanas, H., Rorden, C., Hendrickson, T., Dayan, M., Houlihan, S., Kent, J., Strauss, T., Lee, J., To, I., Markiewicz, C., Lukas, D., Butler, E., Thompson, T., Termenon, M., Smith, D., Macdonald, A., & Kennedy, D. (2024). HeuDiConv — flexible DICOM conversion into structured directory layouts. Journal of Open Source Software. 10.21105/joss.05839

Hutcherson, C. A., Seppala, E. M., & Gross, J. J. (2015). The neural correlates of social connection. Cognitive, Affective, & Behavioral Neuroscience, 15(1), 1–14. 10.3758/s13415-014-0304-9

Jarcho, J. M., Wyngaarden, J. B., Johnston, C. R., Quarmley, M., Smith, D. V., & Cassidy, C. M. (2022). Substance abuse in emerging adults: The role of neuromelanin and ventral striatal response to social and monetary rewards. Brain Sciences, 12(3), 352. 10.3390/brainsci12030352

Jenkinson, M., Beckmann, C. F., Behrens, T. E. J., Woolrich, M. W., & Smith, S. M. (2012). FSL. NeuroImage, 62(2), 782–790. 10.1016/j.neuroimage.2011.09.015

Joyner, K. J., Bowyer, C. B., Yancey, J. R., Venables, N. C., Foell, J., Worthy, D. A., Hajcak, G., Bartholow, B. D., & Patrick, C. J. (2019). Blunted Reward Sensitivity and Trait Disinhibition Interact to Predict Substance Use Problems. Clinical Psychological Science, 7(5), 1109–1124. 10.1177/2167702619838480

Kahneman, D., & Tversky, A. (1979). Prospect theory: An analysis of decision under risk. Econometrica, 47(2), 363–391.

Kim, S. H., Yoon, H., Kim, H., & Hamann, S. (2015). Individual differences in sensitivity to reward and punishment and neural activity during reward and avoidance learning. Social Cognitive and Affective Neuroscience, 10(9), 1219–1227. 10.1093/scan/nsv007

Knutson, B., Westdorp, A., Kaiser, E., & Hommer, D. (2000). FMRI visualization of brain activity during a monetary incentive delay task. NeuroImage, 12(1), 20–27. 10.1006/nimg.2000.0593

Kober, H., Mende-Siedlecki, P., Kross, E. F., Weber, J., Mischel, W., Hart, C. L., & Ochsner, K. N. (2010). Prefrontal–striatal pathway underlies cognitive regulation of craving. Proceedings of the National Academy of Sciences, 107(33), 14811–14816. 10.1073/pnas.1007779107

Kwon, S.-J., Turpyn, C. C., Prinstein, M. J., Lindquist, K. A., & Telzer, E. H. (2022). Self-oriented neural circuitry predicts other-oriented adaptive risks in adolescence: A longitudinal study. Social Cognitive and Affective Neuroscience, 17(2), 161–171. 10.1093/scan/nsab076

Lahnakoski, J. M., Glerean, E., Salmi, J., Jääskeläinen, I. P., Sams, M., Hari, R., & Nummenmaa, L. (2012). Naturalistic fMRI mapping reveals superior temporal sulcus as the hub for the distributed brain network for social perception. Frontiers in human neuroscience, 6, 233. 10.3389/fnhum.2012.00233

Lockwood, P. L., Apps, M. A. J., & Chang, S. W. C. (2020). Is There a ‘Social’ Brain? Implementations and Algorithms. Trends in Cognitive Sciences, 24(10), 802–813. 10.1016/j.tics.2020.06.011

Luijten, M., Schellekens, A. F., Kühn, S., Machielse, M. W. J., & Sescousse, G. (2017). Disruption of Reward Processing in Addiction: An Image-Based Meta-analysis of Functional Magnetic Resonance Imaging Studies. JAMA Psychiatry, 74(4), 387–398. 10.1001/jamapsychiatry.2016.3084

Maleki, N., Sawyer, K. S., Levy, S., Harris, G. J., & Oscar-Berman, M. (2022). Intrinsic brain functional connectivity patterns in alcohol use disorder. Brain Communications, 4(6), fcac290. 10.1093/braincomms/fcac290

Markiewicz, C. J., Gorgolewski, K. J., Feingold, F., Blair, R., Halchenko, Y. O., Miller, E., Hardcastle, N., Wexler, J., Esteban, O., & Goncavles, M. (2021). The OpenNeuro resource for sharing of neuroscience data. eLife, 10, e71774.

Mars, R. B., Neubert, F.-X., Noonan, M., Sallet, J., Toni, I., & Rushworth, M. (2012). On the relationship between the “default mode network” and the “social brain.” Frontiers in Human Neuroscience, 6. https://www.frontiersin.org/articles/10.3389/fnhum.2012.00189

Mars, R. B., Sallet, J., Schüffelgen, U., Jbabdi, S., Toni, I., & Rushworth, M. F. S. (2012a). Connectivity-based subdivisions of the human right “temporoparietal junction area”: Evidence for different areas participating in different cortical networks. Cerebral Cortex (New York, N.Y.: 1991), 22(8), 1894–1903. 10.1093/cercor/bhr268

Mars, R. B., Sallet, J., Schüffelgen, U., Jbabdi, S., Toni, I., & Rushworth, M. F. S. (2012b). Connectivity-based subdivisions of the human right “temporoparietal junction area”: Evidence for different areas participating in different cortical networks. Cerebral Cortex (New York, N.Y.: 1991), 22(8), 1894–1903. 10.1093/cercor/bhr268

McLaren, D. G., Ries, M. L., Xu, G., & Johnson, S. C. (2012). A generalized form of context-dependent psychophysiological interactions (gPPI): A comparison to standard approaches. NeuroImage, 61(4), 1277–1286. 10.1016/j.neuroimage.2012.03.068

Middleton, F. A., & Strick, P. L. (2000). Basal ganglia and cerebellar loops: Motor and cognitive circuits. Brain Research Reviews, 31(2), 236–250. 10.1016/S0165-0173(99)00040-5

Nagy, G. A., Cernasov, P., Pisoni, A., Walsh, E., Dichter, G. S., & Smoski, M. J. (2020). Reward Network Modulation as a Mechanism of Change in Behavioral Activation. Behavior Modification, 44(2), 186–213. 10.1177/0145445518805682

Narr, R. K., Allen, J. P., Tan, J. S., & Loeb, E. L. (2019). Close Friendship Strength and Broader Peer Group Desirability as Differential Predictors of Adult Mental Health. Child Development, 90(1), 298–313. 10.1111/cdev.12905

Nelson, B. D., Perlman, G., Klein, D. N., Kotov, R., & Hajcak, G. (2016). Blunted Neural Response to Rewards as a Prospective Predictor of the Development of Depression in Adolescent Girls. American Journal of Psychiatry, 173(12), 1223–1230. 10.1176/appi.ajp.2016.15121524

Nusslock, R., Abramson, L. Y., Harmon-Jones, E., Alloy, L. B., & Hogan, M. E. (2007). A goal-striving life event and the onset of hypomanic and depressive episodes and symptoms: Perspective from the Behavioral Approach System (BAS) dysregulation theory. Journal of Abnormal Psychology, 116(1), 105–115. 10.1037/0021-843X.116.1.105

Nusslock, R., & Alloy, L. B. (2017). Reward processing and mood-related symptoms: An RDoC and translational neuroscience perspective. Journal of Affective Disorders, 216, 3–16. 10.1016/j.jad.2017.02.001

Nusslock, R., Almeida, J. R., Forbes, E. E., Versace, A., Frank, E., LaBarbara, E. J., Klein, C. R., & Phillips, M. L. (2012). Waiting to win: Elevated striatal and orbitofrontal cortical activity during reward anticipation in euthymic bipolar disorder adults: Nusslock et al. Bipolar Disorders, 14(3), 249–260. 10.1111/j.1399-5618.2012.01012.x

O’Brien, L., Albert, D., Chein, J., & Steinberg, L. (2011). Adolescents Prefer More Immediate Rewards When in the Presence of their Peers: PEERS AND IMMEDIATE REWARDS. Journal of Research on Adolescence, 21(4), 747–753. 10.1111/j.1532-7795.2011.00738.x

O’Doherty, J. P., Cockburn, J., & Pauli, W. M. (2017). Learning, Reward, and Decision Making. Annual Review of Psychology, 68(1), 73–100. 10.1146/annurev-psych-010416-044216

O’Reilly, J. X., Woolrich, M. W., Behrens, T. E. J., Smith, S. M., & Johansen-Berg, H. (2012). Tools of the trade: Psychophysiological interactions and functional connectivity. Social Cognitive and Affective Neuroscience, 7(5), 604–609. 10.1093/scan/nss055

Park, S. Q., Kahnt, T., Dogan, A., Strang, S., Fehr, E., & Tobler, P. N. (2017). A neural link between generosity and happiness. Nature Communications, 8(1), 15964. 10.1038/ncomms15964

Parkinson, C., Kleinbaum, A. M., & Wheatley, T. (2018). Similar neural responses predict friendship. Nature Communications, 9(1), 332. 10.1038/s41467-017-02722-7

Peirce, J., Gray, J. R., Simpson, S., MacAskill, M., Höchenberger, R., Sogo, H., Kastman, E., & Lindeløv, J. K. (2019). PsychoPy2: Experiments in behavior made easy. Behavior Research Methods, 51(1), 195–203. 10.3758/s13428-018-01193-y

Piccirillo, M. L., Lim, M. H., Fernandez, K. A., Pasch, L. A., & Rodebaugh, T. L. (2021). Social Anxiety Disorder and Social Support Behavior in Friendships. Behavior Therapy, 52(3), 720–733. 10.1016/j.beth.2020.09.003

Powers, K. E., Schaefer, L., Figner, B., & Somerville, L. H. (2022). Effects of peer observation on risky decision-making in adolescence: A meta-analytic review. Psychological Bulletin, 148(11–12), 783–812. 10.1037/bul0000382

Rorden, C. (2025). MRIcroGL: voxel-based visualization for neuroimaging. Nature methods. doi:10.1038/s41592-025-02763-7

Rubin, M. (2021). When to adjust alpha during multiple testing: a consideration of disjunction, conjunction, and individual testing. Synthese, 199(3), 10969–11000. 10.1007/s11229-021-03276-4

Salamone, J. D., Correa, M., Yang, J.-H., Rotolo, R., & Presby, R. E. (2018). Dopamine, effort-based choice, and behavioral economics. Frontiers in Behavioral Neuroscience, 12, 52. 10.3389/fnbeh.2018.00052

Santiesteban, I., Kaur, S., Bird, G., & Catmur, C. (2017). Attentional processes, not implicit mentalizing, mediate performance in a perspective-taking task: Evidence from stimulation of the temporoparietal junction. NeuroImage, 155. 10.1016/j.neuroimage.2017.04.055

Saunders, J. B., Aasland, O. G., Babor, T. F., de la Fuente, J. R., & Grant, M. (1993). Development of the Alcohol Use Disorders Identification Test (AUDIT): WHO Collaborative Project on Early Detection of Persons with Harmful Alcohol Consumption--II. Addiction (Abingdon, England), 88(6), 791–804. 10.1111/j.1360-0443.1993.tb02093.x

Saxe, R., & Kanwisher, N. (2013). People thinking about thinking people: the role of the temporo-parietal junction in “theory of mind”. In Social neuroscience (pp. 171–182). Psychology Press. 10.4324/9780203496190

Sazhin, D., Frazier, A. M., Haynes, C. R., Johnston, C. R., Chat, I. K.-Y., Dennison, J. B., Bart, C. P., McCloskey, M. E., Chein, J. M., Fareri, D. S., Alloy, L. B., Jarcho, J. M., & Smith, D. V. (2020). The Role of Social Reward and Corticostriatal Connectivity in Substance Use. Journal of Psychiatry and Brain Science, 5, e200024. 10.20900/jpbs.20200024

Sazhin, D., Wyngaarden, J. B., Dennison, J. B., Zaff, O., Fareri, D., McCloskey, M. S., Alloy, L. B., Jarcho, J. M., & Smith, D. V. (2024). Trait reward sensitivity modulates connectivity with the temporoparietal junction and Anterior Insula during strategic decision making. Biol Psychol, 192, 108857. 10.1016/j.biopsycho.2024.108857

Schultz, R. T., Grelotti, D. J., Klin, A., Kleinman, J., Van Der Gaag, C., Marois, R., & Skudlarski, P. (2003). The role of the fusiform face area in social cognition: Implications for the pathobiology of autism. Philosophical Transactions of the Royal Society of London. Series B: Biological Sciences, 358(1430), 415–427. 10.1098/rstb.2002.1208

Scult, M. A., Knodt, A. R., Hanson, J. L., Ryoo, M., Adcock, R. A., Hariri, A. R., & Strauman, T. J. (2016). Individual differences in regulatory focus predict neural response to reward. Social Neuroscience, 12(4), 419–429. 10.1080/17470919.2016.1178170

Senna, S., Schwab, B., Melo, H. M., Diaz, A. P., & Schwarzbold, M. L. (2022). Social cognition and suicide-related behaviors in depression: A cross-sectional, exploratory study. Brazilian Journal of Psychiatry, 44(6), 639–643. 10.47626/1516-4446-2021-2407

Sequeira, S. L., Silk, J. S., Ladouceur, C. D., Hanson, J. L., Ryan, N. D., Morgan, J. K., McMakin, D. L., Kendall, P. C., Dahl, R. E., & Forbes, E. E. (2021). Association of Neural Reward Circuitry Function With Response to Psychotherapy in Youths With Anxiety Disorders. American Journal of Psychiatry, 178(4), 343–351. 10.1176/appi.ajp.2020.20010094

Shadur, J. M., & Hussong, A. M. (2014). Friendship intimacy, close friend drug use, and self-medication in adolescence. Journal of Social and Personal Relationships, 31(8). 10.1177/0265407513516889

Smith, A. R., Steinberg, L., Strang, N., & Chein, J. (2015). Age differences in the impact of peers on adolescents’ and adults’ neural response to reward. Developmental Cognitive Neuroscience, 11, 75–82. 10.1016/j.dcn.2014.08.010

Smith, D. V., & Delgado, M. R. (2015). Reward Processing. In A. W. Toga (Ed.), Brain Mapping (pp. 361–366). Waltham: Academic Press.

Smith, D. V., & Delgado, M. R. (2017). Meta-analysis of psychophysiological interactions: Revisiting cluster-level thresholding and sample sizes. Human Brain Mapping, 38(1), 588–591. 10.1002/hbm.23354

Smith, D. V., Gseir, M., Speer, M. E., & Delgado, M. R. (2016). Toward a cumulative science of functional integration: A meta-analysis of psychophysiological interactions: Meta-Analysis of Psychophysiological Interactions. Human Brain Mapping, 37(8), 2904–2917. 10.1002/hbm.23216

Smith, D. V., Wyngaarden, J., Sharp, C. J., Sazhin, D., Zaff, O., Fareri, D., & Jarcho, J. (2024). An fMRI dataset of social and nonsocial reward processing in young adults. Data in Brief, 53, 110197. 10.1016/j.dib.2024.110197

Smith, S. M. (2002). Fast robust automated brain extraction. Human Brain Mapping, 17(3), 143–155. 10.1002/hbm.10062

Smith, S. M., Jenkinson, M., Woolrich, M. W., Beckmann, C. F., Behrens, T. E. J., Johansen-Berg, H., Bannister, P. R., De Luca, M., Drobnjak, I., Flitney, D. E., Niazy, R. K., Saunders, J., Vickers, J., Zhang, Y., De Stefano, N., Brady, J. M., & Matthews, P. M. (2004). Advances in functional and structural MR image analysis and implementation as FSL. NeuroImage, 23, S208–S219. 10.1016/j.neuroimage.2004.07.051

Squeglia, L. M., Schweinsburg, A. D., Pulido, C., & Tapert, S. F. (2011). Adolescent Binge Drinking Linked to Abnormal Spatial Working Memory Brain Activation: Differential Gender Effects: ADOLESCENT BINGE DRINKING. Alcoholism: Clinical and Experimental Research, 35(10), 1831–1841. 10.1111/j.1530-0277.2011.01527.x

Strickland, J. C., & Smith, M. A. (2014). The effects of social contact on drug use: Behavioral mechanisms controlling drug intake. Experimental and Clinical Psychopharmacology, 22(1). 10.1037/a0034669

Szanto, K., Dombrovski, A. Y., Sahakian, B. J., Mulsant, B. H., Houck, P. R., Reynolds, C. F., & Clark, L. (2012). Social emotion recognition, social functioning, and attempted suicide in late-life depression. The American Journal of Geriatric Psychiatry, 20(3), 257–265. 10.1097/JGP.0b013e31820eea0c

Telzer, E. H., Fuligni, A. J., Lieberman, M. D., & Galván, A. (2013). Ventral striatum activation to prosocial rewards predicts longitudinal declines in adolescent risk taking. Developmental Cognitive Neuroscience, 3, 45–52. 10.1016/j.dcn.2012.08.004

Torrubia, R., Ávila, C., Moltó, J., & Caseras, X. (2001). The Sensitivity to Punishment and Sensitivity to Reward Questionnaire (SPSRQ) as a measure of Gray’s anxiety and impulsivity dimensions. Personality and Individual Differences, 31(6), 837–862. 10.1016/S0191-8869(00)00183-5

Torrubia, R., & Tobeña, A. (1984). A scale for the assessment of ‘susceptibility to punishment’ as a measure of anxiety: Preliminary results. Personality and Individual Differences, 5(3), 371–375. 10.1016/0191-8869(84)90078-3

Tusche, A., Böckler, A., Kanske, P., Trautwein, F.-M., & Singer, T. (2016). Decoding the Charitable Brain: Empathy, Perspective Taking, and Attention Shifts Differentially Predict Altruistic Giving. Journal of Neuroscience, 36(17), 4719–4732. 10.1523/JNEUROSCI.3392-15.2016

Tziortzi, A. C., Searle, G. E., Tzimopoulou, S., Salinas, C., Beaver, J. D., Jenkinson, M., Laruelle, M., Rabiner, E. A., & Gunn, R. N. (2011). Imaging dopamine receptors in humans with [11C]-(+)-PHNO: Dissection of D3 signal and anatomy. NeuroImage, 54(1), 264–277. 10.1016/j.neuroimage.2010.06.044

Volkow, N. D., Baler, R. D., & Goldstein, R. Z. (2011). Addiction: Pulling at the Neural Threads of Social Behaviors. Neuron, 69(4), 599–602. 10.1016/j.neuron.2011.01.027

Volkow, N. D., Wang, G.-J., Fowler, J. S., Tomasi, D., Telang, F., & Baler, R. (2010). Addiction: Decreased reward sensitivity and increased expectation sensitivity conspire to overwhelm the brain’s control circuit. BioEssays, 32(9), 748–755. 10.1002/bies.201000042

Winkler, A. M., Ridgway, G. R., Webster, M. A., Smith, S. M., & Nichols, T. E. (2014). Permutation inference for the general linear model. NeuroImage, 92, 381–397. 10.1016/j.neuroimage.2014.01.060

Woolrich, M. W., Ripley, B. D., Brady, M., & Smith, S. M. (2001). Temporal Autocorrelation in Univariate Linear Modeling of FMRI Data. NeuroImage, 14(6), 1370–1386. 10.1006/nimg.2001.0931

Wyngaarden, J. B., Johnston, C. R., Sazhin, D., Dennison, J. B., Zaff, O., Fareri, D., McCloskey, M., Alloy, L. B., Smith, D. V., & Jarcho, J. M. (2024). Corticostriatal responses to social reward are linked to trait reward sensitivity and subclinical substance use in young adults. Soc Cogn Affect Neurosci, 19(1). 10.1093/scan/nsae033

Wyngaarden, J. B., Nambiar, A., Dennison, J., Alloy, L. B., Fareri, D. S., Jarcho, J. M., & Smith, D. V. (2025). Trait reward sensitivity and behavioral motivation shape connectivity between the default mode network and the striatum during reward anticipation. bioRxiv, 2025.2004.2017.649386. 10.1101/2025.04.17.649386

Worsley, K. J. (2001). Statistical analysis of activation images. In P. Jezzard, P. M. Matthews, & S. M. Smith (Eds.), Functional Magnetic Resonance Imaging (1st ed., pp. 251–270). Oxford University PressOxford. 10.1093/acprof:oso/9780192630711.003.0014

Young, L., Cushman, F., Hauser, M., & Saxe, R. (2007). The neural basis of the interaction between theory of mind and moral judgment. Proceedings of the National Academy of Sciences, 104(20), 8235–8240. 10.1073/pnas.0701408104

Zahn, R., Moll, J., Krueger, F., Huey, E. D., Garrido, G., & Grafman, J. (2007). Social concepts are represented in the superior anterior temporal cortex. Proceedings of the National Academy of Sciences, 104(15), 6430–6435. 10.1073/pnas.0607061104

Zhang, F., You, Z., Fan, C., Gao, C., Cohen, R., Hsueh, Y., & Zhou, Z. (2014). Friendship quality, social preference, proximity prestige, and self-perceived social competence: Interactive influences on children’s loneliness. Journal of School Psychology, 52(5), 511–526. 10.1016/j.jsp.2014.06.001

